# Long-read single-cell RNA sequencing uncovers cell-type specific transcript regulation in COVID-19

**DOI:** 10.1101/2025.09.23.676511

**Authors:** Kristin Köhler, Lorna Morris, Agata Rakszewska, Robert Lorenz Chua, Maciej Balura, Katharina Jechow, Jennifer Loske, Roland Eils, Irina Lehmann, Peter N. Robinson, Christian Conrad

## Abstract

SARS-CoV-2 infection leads to extensive host transcriptomic changes, but the role of alternative splicing in shaping the immune response remains underexplored. Here, we present the first application of long-read single-cell RNA sequencing on nasopharyngeal swabs from COVID-19 patients and healthy controls to resolve transcript-level changes across cell types. Our analysis identified major epithelial cell types and pronounced immune infiltration, with cell-type annotations concordant with those from short-read data. By enabling isoform-level resolution, our nanopore sequencing approach revealed cell-type specific alternative splicing, undetectable with short-read sequencing. For example, although gene-level expression of the key immune and apoptosis regulators, *IFNAR2* and *FAIM*, did not differ between COVID-19 patients and healthy controls we identified marked shifts in isoform usage. Between moderate and critical cases, we observed cell-type specific differential transcript usage in the T cell signaling kinase *FYN* and the immune-regulatory transcription factor *IRF2*. As some of these splicing alterations yield functionally distinct isoforms, we hypothesize that alternative splicing modulates immune signaling and apoptosis, fine-tuning the host response to SARS-CoV-2 infection. Our study demonstrates the unique power of long-read single-cell transcriptomics to uncover isoform-resolved regulatory changes, offering novel insights into the role of alternative splicing in shaping immune responses to viral infections.

## Introduction

The global impact of the coronavirus disease 2019 (COVID-19) pandemic has underscored the importance of understanding the molecular intricacies of viral infections. COVID-19, caused by the severe acute respiratory syndrome coronavirus-2 (SARS-CoV-2), has presented immense challenges to public health, necessitating a comprehensive exploration of the molecular mechanisms following infection. SARS-CoV-2, which primarily infects cells of the respiratory system, is associated with extremely heterogeneous symptoms [1,2]. While most SARS-CoV-2 infections cause mild respiratory symptoms, approximately 5% of patients develop a critical disease including acute respiratory distress syndrome (ARDS), septic shock or multi-organ failure, all associated with high mortality [3].

Single-cell RNA sequencing (scRNA-seq) has been extensively employed to investigate the cellular and molecular mechanisms underlying COVID-19 [2,4–7]. It has revealed shared gene expression programs in nasal, lung and gut tissues, including genes involved in viral entry, key immune functions, and epithelial–macrophage crosstalk [6]. Additionally, scRNA-seq studies have shown increased expression of pro-inflammatory cytokines in patients with critical COVID-19 [2,7]. However, all scRNA-seq studies of viral infections to date have relied on short-read sequencing technologies, which limits isoform-level insights due to their 3′ or 5′ transcript capture bias [8].

Alternative splicing of transcripts provides an additional regulatory layer to cellular function, in part by increasing protein diversity, often in a cell type-specific manner [9]. Modulations in splicing influence physiological processes including immune responses and have been implicated in various diseases including viral infections [10–13]. The SARS-CoV-2 NSP16 protein has been shown to bind to the spliceosomal U1 and U2 small nuclear RNAs and act to suppress global mRNA splicing during SARS-CoV-2 infection [14]. Moreover, dysregulation of alternative splicing has been observed in COVID-19 patients, distinct alternative splicing patterns of genes related to Janus kinase (JAK) signaling pathway and Toll-like receptor 4 (TLR4) with four different Sars-CoV-2 variants (Alpha, Beta, Gamma, and Omicron), as well as healthy controls [11]. The betacoronoviruses, SARS-CoV, SARS-CoV-2 and MERS exhibit differential alternative splicing of genes that affect a diverse set of biological functions, including genes that target a broad range of cellular functions, with a particular impact on translation, RNA processing, and ribosome biogenesis [10].

Long-read sequencing technologies such as Oxford Nanopore (ONT) and PacBio overcome the limitations of short reads in isoform-level analysis by enabling single-cell full-length isoform identification [15–24]. Until recently, the relatively higher error rate of nanopore sequencing necessitated generation of matched short-read data for error-free and efficient cell barcode assignment [16]. However, improvements in nanopore sequencing accuracy and increased long-read sequencing throughputs have enabled the development of methods such as Nanopore’s *wf-single-cell* and BLAZE [25], which can accurately and efficiently identify 10x cell barcodes using only long-read scRNA-seq data.

In this study, we leverage these advances to perform full-length isoform-resolved single-cell RNA sequencing on nasal and nasopharyngeal swabs from COVID-19 patients and healthy controls. We implemented an analysis pipeline to compare the transcriptomes of patients with moderate and critical COVID-19 with those of healthy controls and explored the functional consequences of differential transcript usage across cell types to better understand the role of alternative splicing in COVID-19 pathogenesis. To our knowledge, this is the first single-cell long-read transcriptomic study of a viral infection, providing new insights into how alternative splicing shapes host responses *in vivo*.

## Results

We generated long-read, scRNA-seq data from nasal and nasopharyngeal samples obtained from nine COVID-19 patients and three SARS-CoV-2–negative donors with no symptoms of respiratory tract infection. Among the patients, three were categorized as moderate while six were critical, as classified by World Health Organization (WHO) guidelines (Supplementary Table S1). All samples had been previously sequenced using Illumina short-read sequencing technology and were part of two studies that characterized and compared COVID-19 patients by severity [2] and age group [26].

In this study, we sequenced unfragmented cDNA aliquots from these samples using ONT to obtain full-length transcript information. Long-read sequencing yielded between 41 million and 98 million quality-filtered reads per flow cell, totaling 786 million reads across all samples (Supplementary Fig. S1b). Mean read quality ranged from 15.5 to 17 for samples using V14 chemistry (see Supplementary Figure S1a). On average, 57% of reads displayed the expected structure—including adapters, unique molecular identifiers (UMIs), and cell barcodes; 47% could be confidently assigned to a gene and 28% to a specific transcript (Supplementary Fig. S1b). The *wf-single-cell* analysis pipeline filters out invalid or low-quality reads through a series of steps. Barcode extraction and correction are performed using the 10x Genomics whitelist of valid barcode sequences. With the default pipeline parameters that were employed in this study, barcodes are corrected if they are within an edit distance of 2, while reads with barcode sequences above this threshold are discarded. The fractions of mapped reads we obtained are consistent with those reported in previous studies. [15]. *wf-single-cell* retains only reads that map to annotated transcripts; novel transcripts are excluded. This limitation, together with the lower reference support for rare transcripts, likely contributes to the reduction of retained reads observed in transcriptome mapping compared to genome mapping.

For downstream analysis, gene-level expression data was used for quality control, dimensionality reduction, clustering, and cell type annotation. After filtering low-quality cells, a total of 31,546 cells remained (Supplementary Fig. S1). Across all cells, 190 million UMIs were assigned to genes and 104 million to specific transcripts. In disease samples, cells exhibited a median of 3,346 gene-assigned UMIs and 2,048 transcript-assigned UMIs, whereas in control cells the medians were 3,836 gene-assigned and 1,843 transcript-assigned UMIs per cell (Supplementary Fig. S2a). The lower sequencing depth observed in controls reflects the higher number of cells in these samples (Supplementary Fig. S1d).

Based on the gene-level analysis, we identified six major epithelial and immune cell types (Figure 1b). All previously described epithelial cell types of the upper respiratory tract were identified based on their marker genes (Figure 1c), including basal, secretory, FOXN4^+^, mature ciliated cells and differentiating ciliated cells, as well as ionocytes. Immune cell types comprised macrophages/monocytes, neutrophils, B cells, CD4^+^ and CD8^+^ T cells and mast cells, all expressing respective marker genes (Figure 1c) [2,27].

**Figure 1:**
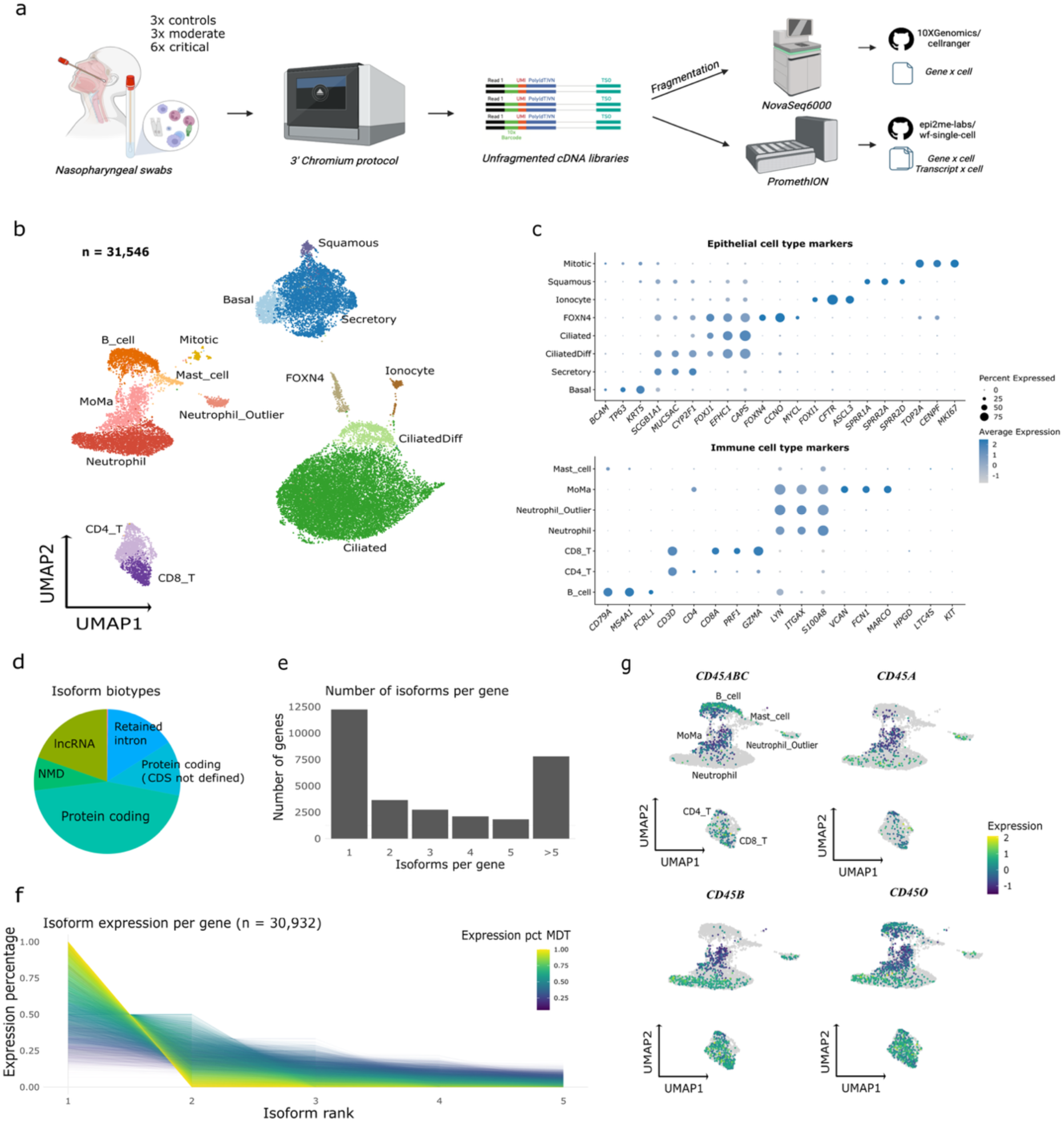
Long-read sequencing identifies major immune and epithelial cell types. **(a)** Overview of single-cell long-read sequencing workflow. Samples were processed using droplet-based single-cell RNA-seq 3’-**g**ene expression assay to obtain cDNA. Libraries for third-generation sequencing were prepared from unfragmented cDNA after depletion of artefactual molecules. Sequencing was carried out on the PromethION instrument, followed by data processing with wf-single-cell (see Materials and Methods). (b) Gene expression-based UMAP embedding of cells (n = 31,546) identified from long-read sequencing. (c) Dot plots displaying marker gene expression for epithelial and immune cell populations present in the upper respiratory tract (d) Pie chart depicting the proportions of Ensembl biotype classifications among the identified isoforms (e) Bar plot showing the number of isoforms identified per gene (f) Isoforms per gene ranked by their expression percentage: each line represents one gene showing the expression values of its most expressed isoforms on the y-axis. Line colors denote the proportional expression of the gene’s most dominant transcript (MDT). (g) Expression of known CD45 isoforms (encoded by *PTPRC*) O, A, B and ABC projected onto UMAP embedded immune cell populations.

Consistent with previous reports, cell type distributions varied considerably between samples and conditions [2]. Samples from COVID-19 patients were dominated by immune cells, with a significantly higher immune-to-epithelial cell ratio compared to controls (Fisher’s exact test, p < 2.2 × 10⁻¹⁶, OR = 114.0, 95% CI: 103.5–125.1), consistent with immune cell recruitment to the primary site of infection (Supplementary Fig. S2c). Across all cells, we identified 32,460 genes and 128,027 distinct transcripts. For most genes, only a single isoform was detected (Figure 1e). Since sequencing depth influences the number of detected isoforms (Supplementary Fig. S2d), it is likely that additional low-abundance isoforms remain undetected. Most genes expressed a single dominant isoform, whereas some genes exhibited more balanced expression across multiple isoforms (Figure 1f). According to GENCODE, most identified transcripts were classified as protein-coding, with 45% containing a known coding sequence and 12% containing an unknown coding sequence (Figure 1d). Approximately 19% are annotated as long non-coding RNAs (lncRNAs), while 16% contain retained introns. Transcripts containing retained introns could represent splicing intermediates or transcripts that regulate gene expression, for example, by competing with fully spliced mRNAs for translation. Additionally, 7% of transcripts are known nonsense-mediated decay (NMD) targets, leading to their degradation due to premature stop codons. These transcriptomic patterns are largely consistent between controls, moderate, and critical COVID-19 patients (Supplementary Fig. S3).

A well-known example of differential transcript usage in immune cell types is the *PTPRC* (CD45 antigen) gene [28]. CD45 is a key immune cell marker with distinct isoform expression patterns among different subpopulations of immune cells. To validate our transcript-level signals, we assessed differential expression of *PTPRC* (CD45) isoforms across immune cell subpopulations and projected isoform expression onto the gene-level Uniform Manifold Approximation and Projection (UMAP). As previously reported, the longest isoform, *CD45ABC*, serves as a pan-B-cell marker [29], and is strongly upregulated in our B cell population (two-sided Wilcoxon rank-sum test, avg log₂ FC = 2.27, p = 1.3 × 10⁻^128^; Fig. 1d). In T cells, the naïve marker *CD45RA* shows only a modest increase in expression (avg log₂ FC = 0.14, p = 0.35), whereas the memory/effector isoform *CD45RO* is significantly upregulated (avg log₂ FC = 1.29, p = 1.1 × 10^-35^) [30], consistent with an activated phenotype. The regulatory-associated *CD45RB* is also upregulated in both CD4⁺ and CD8⁺ T-cell subsets (avg log₂ FC = 1.47, p = 2.9 × 10^-54^). Monocytes and macrophages express all CD45 isoforms at lower levels compared to other immune cells (*CD45ABC*: avg log₂ FC = −0.51, p= 6.4 × 10^-13^; *CD45A*: −0.38, p = 6.0 × 10^-32^; *CD45B*: −1.48, p = 3.2 x 10^-7^; *CD45O*: −0.88, p = 4.3 × 10^-12^).

### Cell-type annotations from short- and long-read scRNA-seq data exhibit high concordance

Next, we compared the ability of long- and short-read scRNA-seq data to cluster cells and annotate cell types. Both approaches showed qualitative similarity in UMAP visualizations of identified cell populations and overall cell type distributions (Fig. 2a, b). Projecting short-read based annotations onto the long-read UMAP showed that most cells were assigned the same types in both modalities (Fig. 2c). To quantify this, we performed label transfer from the short-read to the long-read dataset, which yielded consistently high transfer scores, indicating high concordance in cell-type identities between the two sequencing approaches (Fig. 2d).

**Figure 2:**
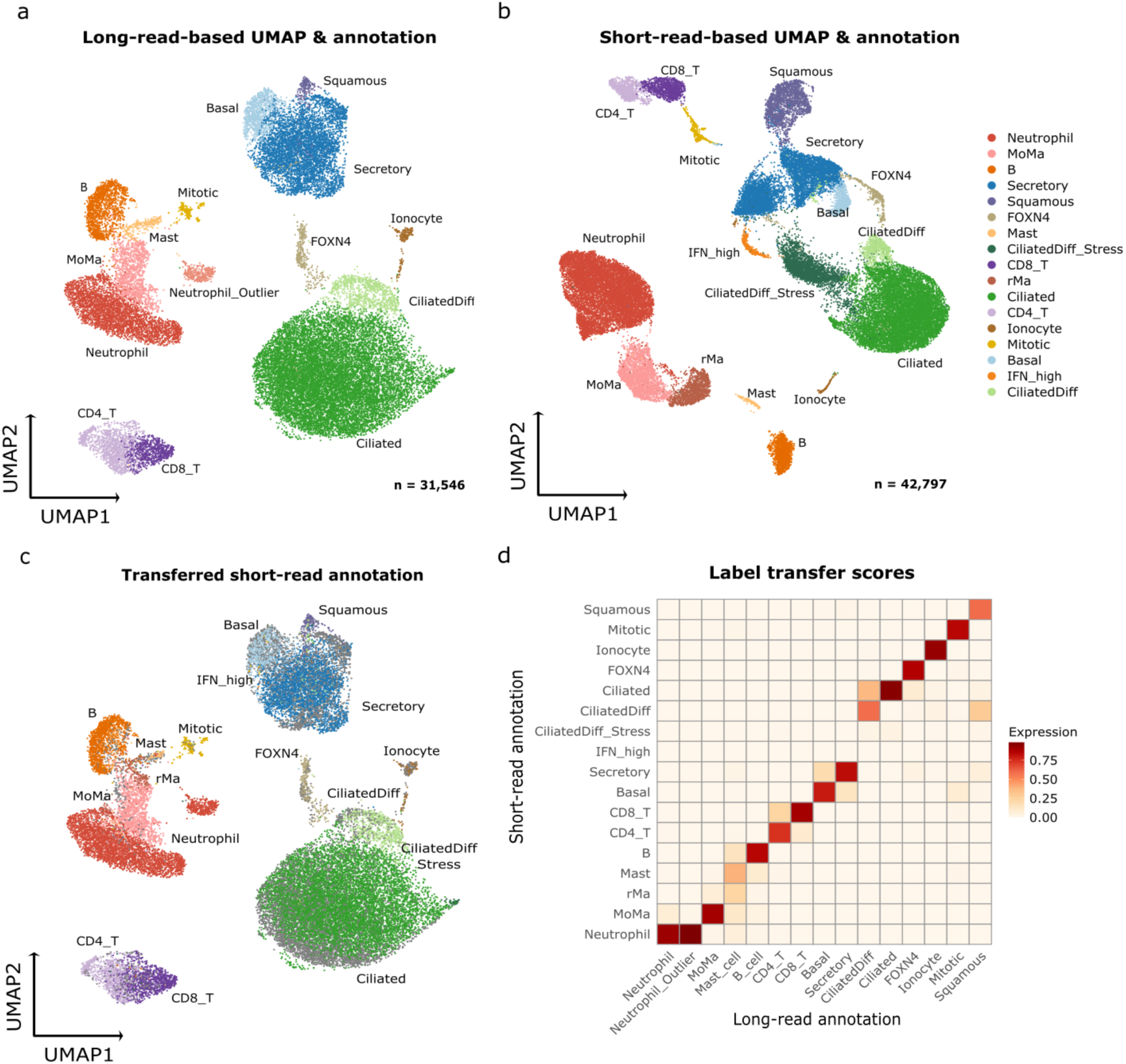
Comparison of short- and long-read RNA-seq. (a) Gene expression-based UMAP embedding of cells identified from long-read RNA-seq (n = 31,546) (b) Gene expression-based UMAP embedding of cells identified from short-read sequencing (n = 42,797) (c) Annotations obtained from short-read dataset transferred onto the long-read defined UMAP embedding. (d) Heatmap displaying the average label transfer scores obtained from transferring annotations from the short-read cell embeddings to cells identified from long-read sequencing.

However, we observed differences in total cell numbers, with 31,546 cells identified in the long-read dataset and 42,797 in the short-read dataset. Cells unique to the short-read dataset exhibited lower complexity, with a median of 489 detected genes and 895 UMIs per cell. This discrepancy can be attributed to the lower sequencing depth of the long-read dataset and results in some of the macrophages, IFN-responsive cells, and a proportion of neutrophils, ciliated, and squamous cells to be detected only in the short-read dataset (Supplementary Fig. S5).

### Differentially expressed genes between COVID-19, controls, and disease severities also show differential transcript usage

We performed differential gene expression (DGE) and differential transcript usage (DTU) analysis between COVID-19 and control samples (Fig 3a), and moderate versus critical samples (Fig 3b), using pseudo-bulked data. For DGE, we used DESeq2 [31], which models gene-level counts using a negative binomial generalized linear model (GLM) and tests for differential expression using the Wald test and Benjamini-Hochberg correction. For DTU, we applied DEXSeq [32], which also uses negative binomial GLMs to model transcript-level counts and identifies differential transcript usage by testing for significant condition– transcript interactions using a likelihood ratio test (LRT), followed by Benjamini-Hochberg p-value correction. Pseudobulk samples were obtained by aggregating gene or transcript expression counts across all cells of the same annotated type within each individual sample. In COVID-19 vs. control the highest number of significant DEGs (> 750 genes) were detected in ciliated and secretory cells and much lower numbers (10 or fewer) were detected in squamous, FOXN4^+^ and plasma B cells, whilst the other cell types had between 50 and 300 DEGs (p_adj_ < 0.05, Figure 3a).

**Figure 3:**
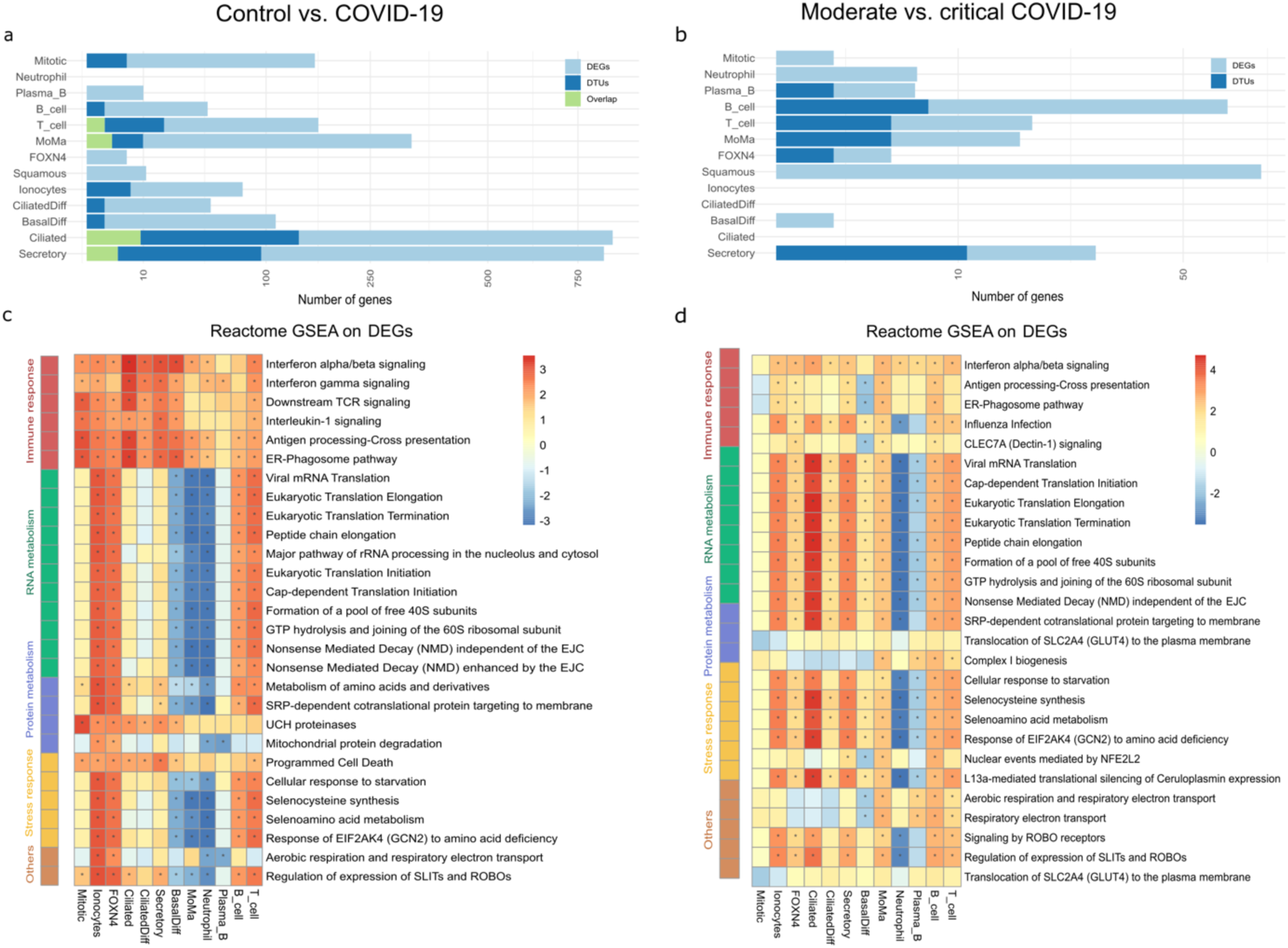
Differential expression analysis of long-read RNA-seq reveals shared and distinct enrichment in biological pathways across cell-types in COVID-19 patients. (a,b) Bar plots displaying number of significant DEGs (Wald test, Benjamini-Hochberg adjusted two-tailed, negative-binomial p_adj_ < 0.05) and DTU genes (Log-Likelihood-Ratio test) identified per cell type for (a) COVID-19 vs control and (b) moderate vs critical COVID-19. (c,d) Heatmaps showing normalized enrichment scores of top 10 enriched Reactome pathways from GSEA across all cell types for DEGs in (c) COVID-19 vs control and (d) moderate vs critical COVID-19.

In comparison, there were considerably fewer genes significantly differentially expressed between moderate and critical disease, with more than 10 occurring only in B cells, T cells, monocytes and macrophages, squamous and secretory cells (Figure 3b). Figure 3a and 3b also show the differentially expressed genes that additionally demonstrated significant DTU (p_adj_ < 0.05) and the DTU genes that are not differentially expressed at the gene-level between the conditions. Fewer significant DTU genes were detected per cell type than DEGs, which may reflect data sparsity and lower transcript counts, or indicate that fewer genes are regulated at the transcript level via mechanisms like alternative splicing, transcription initiation and termination. The highest number of DTU genes were detected in secretory and ciliated cells with 100 and 150 genes identified, respectively. DEGs with concurrent differential transcript usage were only detected when comparing COVID-19 patients to controls, specifically in ciliated, secretory, MoMa (monocytes and macrophages), and T cells—cell types that also showed the highest overall number of differentially expressed genes compared to other cell types.

The majority of both upregulated and downregulated transcripts in COVID-19 were protein-coding. Transcripts annotated as nonsense-mediated decay (NMD) and lncRNAs were more frequently upregulated in COVID-19, while transcripts with retained introns were more often downregulated. (Supplementary Fig. S6).

### Gene Set Enrichment Analysis reveals genes in enriched pathways are regulated at both the gene and the transcript level

To investigate the functional consequences of gene expression differences between COVID-19 of different severities and healthy controls we performed Gene Set Enrichment Analysis (GSEA) [33]. Genes were ranked by the DESeq2 Wald test statistic, which reflects both the magnitude of differential expression and its variability. The top enriched Reactome pathways fell into four broad categories: Immune response, RNA metabolism, protein metabolism and stress response (Figure 3c and 3d). Enriched pathways were identified even in cell types with few or no significant DEGs, as GSEA considers the full ranked list of genes rather than only those below the significance threshold (p_adj_ < 0.05).

In patients with COVID-19, there was a significant enrichment of immune response pathways in most cell types, with the highest enrichment score observed in ciliated cells, for ‘IFN alpha / beta signaling’, ‘IFNγ signaling’, ‘Interleukin I signaling’, ‘Downstream TCR signaling’ and ‘ER Phagosome Pathway’ (Fig 3c). Although immune response pathways were also enriched when comparing gene expression from moderate with critical patients, only ‘IFN alpha / beta signaling’ was observed as enriched across all cell types, and in general enrichment scores were lower indicating that gene expression differences were less pronounced (Figure 3d). Moreover, the Reactome pathway ‘Influenza infection’ was negatively enriched in neutrophils and positively enriched in most other cell types, corresponding to the downregulation of ribosomal protein large and small subunit genes (RPL/RPS) in neutrophils and their upregulation in other cell types.

Several Human Leukocyte Antigen (HLA) genes, including *HLA-A*, *HLA-B*, *HLA-C* and *HLA-E*, were significantly upregulated in COVID-19 vs controls. These genes also consistently appeared among the leading-edge genes in inflammatory response pathways, suggesting they are common drivers of these enrichments. While the immune response pathways were enriched in most cell types, the involvement of RNA and protein metabolism pathways showed greater cell-type specificity. The RNA metabolism pathways include translation elongation and termination and nonsense mediated RNA decay, of which the leading-edge genes in these enriched gene sets contained predominantly RPL/RPS genes.

Pathways relating to programmed cell death were enriched across all epithelial cells and T cells, with the strongest signal observed in secretory cells (Figure 3c). In these cells, the most significantly differentially expressed genes were *UBC* and *CASP4*. In contrast, in ciliated cells, several genes encoding components of the 26S proteasome complex (*PSMA*, *PSMB*, *PSMD*, and *PSME*) were differentially expressed in healthy vs COVID-19.

We also performed GSEA using the Hallmark gene sets of the Molecular Signatures Database (MsigDB) [34] and Gene Ontology Biological process gene sets [35] and found similar pathways, for example ‘Hallmark Interferon alpha’ and ‘interferon gamma response’ in all cell types enriched for our datasets (Supplementary tables). The Hallmark complement pathway genes were significantly enriched in several immune cell types - monocytes and macrophages, neutrophils and T-cells but not in epithelial cells.

### Isoform switches between COVID-19 and healthy controls occur in key immune response pathways

Next, we focused on key pathways upregulated in COVID-19 and identified several pathway genes with differential transcript usage. Notably, most DTU genes do not show significant changes in overall gene expression levels but rather exhibit shifts in relative transcript usage—suggesting that immune regulation may be fine-tuned through alternative splicing or virus-induced splicing alterations. Although we performed pathway analysis on genes with differential transcript usage, the results were not statistically significant, likely due to the relatively small number of DTU genes compared to differentially expressed genes. To assess whether differential transcript usage was a potential regulatory mechanism in the biological pathways detected from gene-level analysis we mapped genes that had significant differential transcript usage to the most enriched pathway gene sets from the GSEA analysis (Figure 4a, e, g and 5a, e, f).

**Figure 4:**
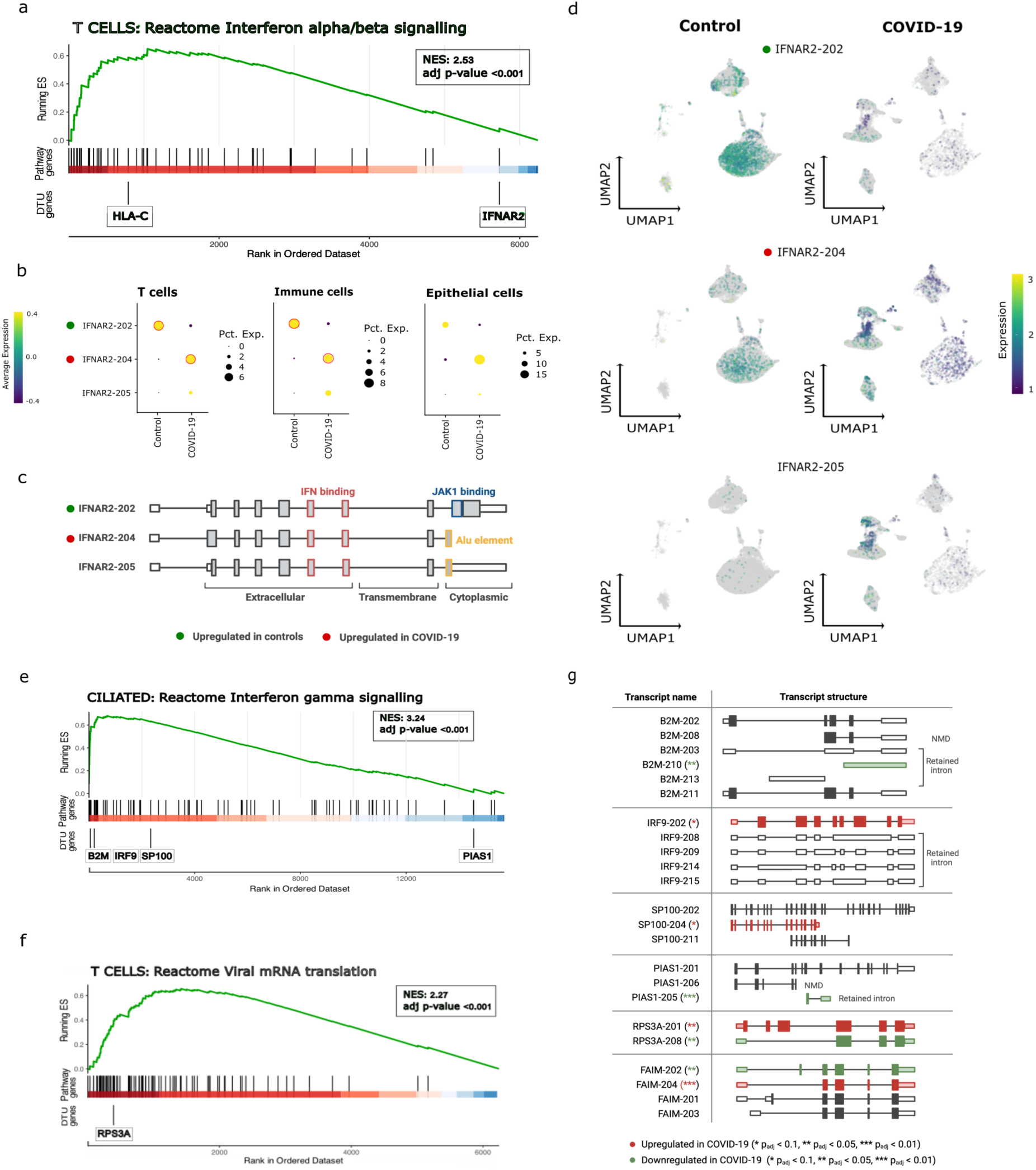
Key genes in the immune response to viral infection in COVID-19 are not only differentially expressed but also regulated at the transcript level. (a) GSEA plot showing genes ranked by the Wald statistic from the differential gene expression (DGE) analysis between control and COVID-19 samples in T cells. The x-axis indicates the position of genes in the ranked list and the track labeled “Pathway genes” indicates the presence of the gene in the ranked list in the *Interferon alpha/beta signaling* pathway. The green line represents the running enrichment score across the ranked gene list. Genes with significant DTU (Benjamini-Hochberg adjusted two-tailed, negative-binomial; *p_adj_* < 0.05) are highlighted in the bottom track. (b) Dot plot showing expression of *IFNAR2* isoforms across T cells, all immune and epithelial cell populations combined. Red circles indicate significant change in transcript usage between conditions. We observe a significant reduction from IFNAR2-202 (L) to IFNAR2-204 (S) from control samples to COVID-19 samples in T cells (log_2_FC = −3.34, p_adj_ = 0.005 for *IFNAR2-202*; log_2_FC = 4.0, p_adj_ = 0.002 for *IFNAR2-204*) and immune cell populations (log_2_FC = −3.2, p_adj_ = 0.003 for *IFNAR2-L*; log_2_FC = 3.3, p_adj_ = 0.003 for *IFNAR2-S*). (c) *IFNAR2* isoform schematics displaying exon structures and associated functional domains (d) Expression of *IFNAR2* isoforms projected onto the UMAP embedded cell populations. (e) GSEA plot showing enrichment of genes involved in the *Interferon gamma signaling* pathway in ciliated cells (genes ranked by the Wald statistic) (f) GSEA plot showing enrichment of genes involved in the viral mRNA translation pathway in T cells (genes ranked by Wald statistic). (g) Genes exhibiting DTU between control and COVID-19 samples. Transcript bars are color-coded according to the direction of expression change. Asterisks denote the statistical significance of DTU (Benjamini-Hochberg adjusted p-values from two-tailed negative binomial LRT testing: * padj < 0.1, ** padj < 0.05, *** padj < 0.01).

A key player in the interferon alpha/beta pathway is *IFNAR2*, the receptor subunit that binds type I interferons and triggers JAK-STAT signaling leading to the transcription of interferon-stimulated genes (ISGs). In our dataset, *IFNAR2* was not significantly differentially expressed in any cell type. Notably, although the Reactome interferon alpha/beta pathway was enriched, *IFNAR2* was ranked near the bottom of the pathway’s gene list, suggesting that *IFNAR2* expression contributed minimally to the observed enrichment (Figure 4a). However, we detected a significant shift in *IFNAR2* transcript usage in COVID-19 patients compared to controls, with a reduction in the canonical *IFNAR2-202* isoform and an increase in *IFNAR2-204* across all immune cells, particularly T cells. A similar pattern was observed in epithelial cells, but it was not consistent across control samples and thus not statistically significant.

GSEA analysis of DEGs between COVID-19 and control samples revealed the Interferon gamma signaling pathway with the highest enrichment score in ciliated cells (NES = 3.24, Figure 3, Figure 4e). This pathway also included four significant DTU genes: *B2M, IRF9, SP100,* and *PIAS1*. While *B2M* shows a significant change in gene expression in COVID-19 (logFC = 1.00, p_adj_ = 0.01), all other genes exhibit changes only in transcript usage (Table 1, Supplementary Fig. S5). In *PIAS1*, *B2M,* and *IRF9*, we observe switches from short transcripts with retained introns in control samples to protein-coding isoforms in COVID-19. In *SP100*, we observe a switch from shorter, protein-coding isoforms lacking multiple 5’ exons to two longer, protein-coding isoforms (Table 1, Supplementary Fig. S6).

In T cells the viral mRNA translation pathway is enriched, driven in part by differential expression of RPL and RPS gene family members between COVID-19 and control samples. Notably, one of these genes, *RPS3A*, exhibits differential transcript usage in T cells (Figure 4g). We observe a switch from the shorter isoform *RPS3A-208* in controls to the longer *RPS3A-201* isoform in COVID-19 (Table 1, Supplementary Fig. S6).

A notable isoform switch was observed in the Fas apoptosis inhibitory molecule (*FAIM*) gene. *FAIM*, a negative regulator of Fas-mediated apoptosis, was not differentially expressed at the gene level and was not annotated in any of the gene sets used in the enrichment analysis. We identify two isoforms of *FAIM* as differentially used in ciliated cells (Figure 4i) with the shorter *FAIM-204 (S)* isoform highly upregulated in COVID-19 and the longer canonical isoform, *FAIM-202 (L)*, solely expressed in our controls (Table 1, Supplementary Fig. S6).

### Isoform switches between moderate and critical COVID-19 occur in key immune response pathways

We employed the same strategy to compare the DTU genes between moderate and critical COVID-19 cases as we used for comparing COVID-19 with healthy controls. However, because there were fewer DEGs and scores for Reactome pathway enrichment were lower than the comparison with healthy controls (Figure 3d), we expanded our analysis to include Hallmark and GO Biological Process gene sets. This switch allowed us to explore a broader and more diverse range of functional categories: Hallmark gene sets are often broader in scope and can capture general biological processes, while GO terms offer more granular coverage across a wider range of annotated functions. Using these gene sets, we identified additional DTU genes within enriched pathways that were not captured using Reactome.

In the top 5 enriched Hallmark pathways in T cells, the Hallmark Complement gene set contains the gene *FYN*, which has differential transcript usage but is not differentially expressed between moderate and critical COVID-19 (Figure 5a). *FYN* is also found in the top enriched pathway, GO Biological Process – ‘intrinsic apoptosis signaling’ pathway. We detected four protein-coding isoforms, *FYN-202, FYN-206, FYN-207, and FYN-229* (Figure 5b,c). The two longer isoforms, *FYN-229, FYN-202*, consist of 10 exons and differ in an alternate exon 5, while *FYN-207* includes exons 5 to 10 and the shortest isoform, *FYN-206* consists of only the terminal two 3 prime exons. Only the longer isoforms, *FYN-229* and *FYN-202* contain the protein kinase domains.

**Figure 5:**
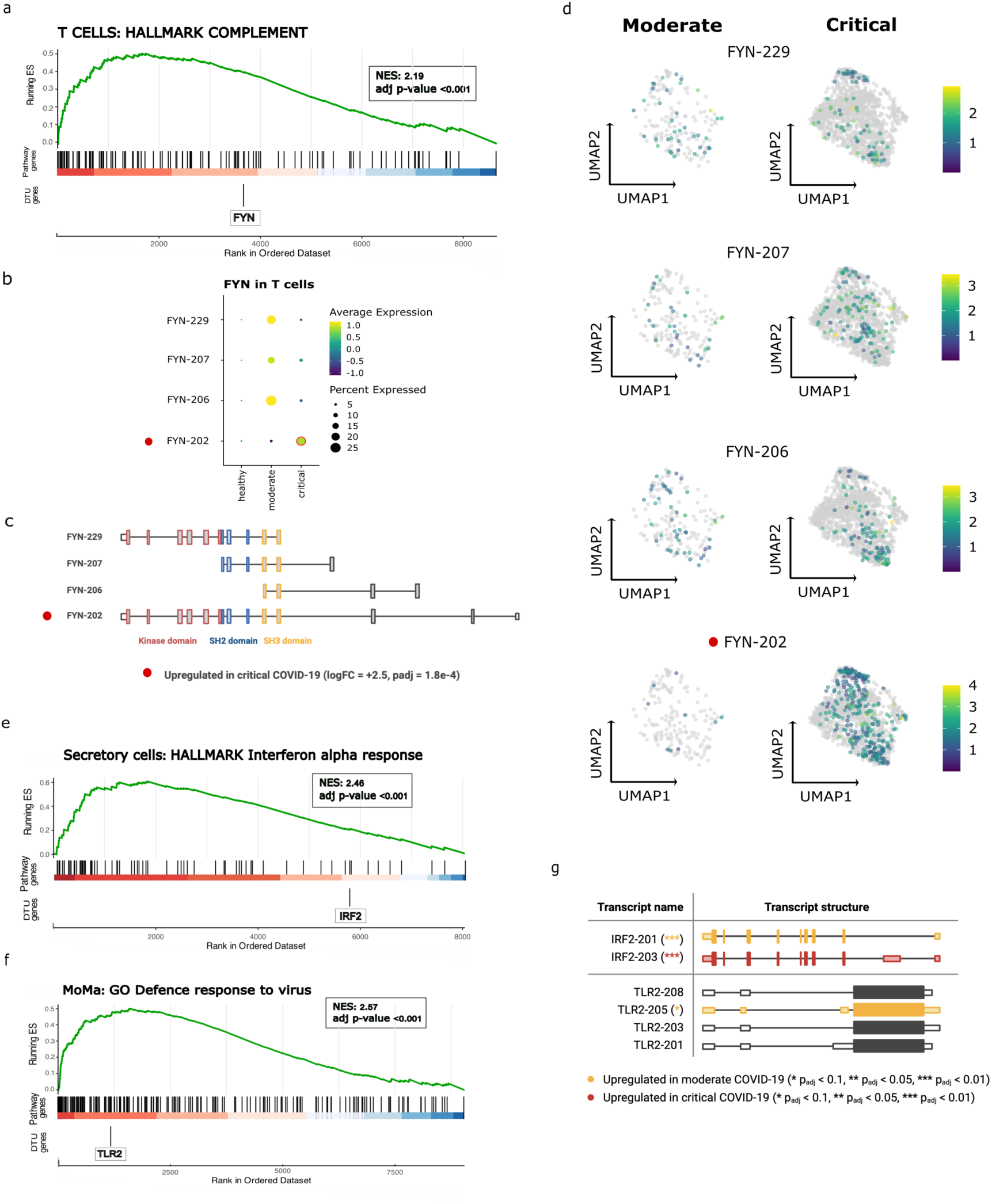
Key genes in the immune response and defense response to virus in critical COVID-19 are not only differentially expressed but also regulated at the transcript level. (a) GSEA plot showing genes ranked by the Wald statistic from the differential gene expression (DGE) analysis between moderate and critical COVID-19 in T cells. Genes that are involved in the *Hallmark Complement* pathway are highlighted in the middle track. The green line represents the running enrichment score across the ranked gene list. Genes with significant DTU (Benjamini-Hochberg adjusted two-tailed, negative-binomial; *p_adj_* < 0.1) are highlighted in the bottom track. (b) Dot plot showing expression of *FYN* isoforms across T cells. Red circles indicate significant change in transcript usage between conditions. We observe a significant increase of FYN-202 usage in critical COVID-19 in T cells (log_2_FC = 2.52, p_adj_ = 1.8 x 10^-4^) (c) FYN isoform schematics displaying exon structures and associated functional domains (d) Expression of FYN isoforms projected onto the UMAP embedded cell populations (e) GSEA plot showing enrichment of genes involved in the *Hallmark: Interferon alpha response* pathway in secretory cells (genes ranked by the Wald statistic) (f) GSEA plot showing enrichment of genes involved in the *GO: Defence response to virus* pathway in T cells (genes ranked by Wald statistic). (g) Genes exhibiting DTU between moderate and critical COVID-19. Transcript bars are color-coded according to the direction of expression change. Asterisks denote the statistical significance of DTU (Benjamini-Hochberg adjusted p-values from two-tailed negative binomial LRT testing: * padj < 0.1, ** padj < 0.05, *** padj < 0.01).

The Hallmark Interferon alpha pathway was enriched in COVID-19 vs control and to a lesser extent in critical vs moderate COVID-19 (Figure 5d), although differential transcript usage of *IRF2* was only detected between moderate and critical COVID-19. *IRF2* was not differentially expressed at the gene-level, but two different isoforms (*IRF2-201* and *IRF2-203*) were detected (Table 2, Supplementary Fig. S6). Both encode proteins of the same length but differ in their 5′ untranslated regions (UTRs). The canonical isoform, *IRF2-201* was more highly expressed in critical COVID-19, while *IRF2-203* was more highly expressed in moderate COVID-19 (Table 2, Supplementary Fig. S6). Another enriched pathway between moderate and critical COVID-19 was *GO: Defense response to virus*. One of its differentially expressed genes, *TLR2*, also exhibited differential transcript usage (Table 2, Supplementary Fig. S6). Four *TLR2* isoforms were detected, each with differing UTRs and a single protein-coding exon.

## Discussion

Using long-read sequencing, we demonstrate that key genes involved in the immune response to viral infection are regulated at the transcript level through mechanisms including alternative splicing, even when gene expression levels show no significant changes. To our knowledge, this is the first single-cell study of isoform level expression in patients with COVID-19.

Profiling more than 30,000 cells with long-read scRNA-seq, we identified and characterized epithelial cell populations in the human nasopharynx and immune cells recruited during COVID-19. We show a high agreement between cell populations identified in short and long-read sequencing data and their marker gene expression. However, due to lower sequencing depth in long-read sequencing, cell populations with physiologically or technically low transcript abundances are underrepresented. While increasing sequencing depth would likely mitigate the underrepresentation of populations with lower transcript abundance, this is more costly with long-read sequencing.

Although low sequencing depth limited detection of some populations, gene set enrichment analysis revealed substantial alterations in immune and stress response pathways, as well as RNA and protein metabolism, with both conserved and distinct regulation patterns across cell types. Notably, among the pathway genes, we identified several genes that are exclusively regulated at the level of splicing/transcript usage, without changes in overall expression.

In COVID-19 compared to healthy donors, we observed a shift in expression, from the canonical *IFNAR2-202* isoform to the truncated *IFNAR2-204* isoform across immune cells, which was most pronounced in T cells. There has been significant interest in understanding the role of *IFNAR2* in COVID-19, with several studies investigating its involvement in disease severity and immune response regulation [38–41]. Reduced *IFNAR2* expression was linked to severe COVID-19 outcomes, and genomic variants in *IFNAR2* have been associated with altered interferon binding and increased mortality risk [41]. Recent studies suggest that differential isoform expression of *IFNAR2* may influence COVID-19 outcome [42]. The canonical *IFNAR2-L (IFNAR-202)* isoform activates JAK-STAT signaling upon binding to type I interferons, leading to the upregulation of ISGs [42]. While the truncated *IFNAR2-S (IFNAR-204)* isoform, lacking the cytoplasmic domain, has been proposed to act as a decoy receptor by binding interferons without activating JAK-STAT signaling [42], it may modulate IFN responses and prevent overactivation of ISGs. However, *IFNAR2-S* knockout in A549 cell lines was shown to enhance the antiviral response to SARS-CoV-2, indicating that *IFNAR2-S* overexpression might inhibit the protective effect of interferons. Our findings, together with the observations from previous cell line experiments, highlight the need to investigate *IFNAR2* isoform dynamics and their functional consequences in longitudinal patient cohorts.

We identified several other DTU genes involved in interferon-γ signaling in ciliated cells, including *B2M*, *IRF9* and *PIAS1*, which shifted from retained-intron to fully spliced, protein-coding transcripts in COVID-19. Notably, the *SP100-204* isoform (Sp100A), known to promote ISG expression through chromatin remodeling [43–45], was significantly upregulated in COVID-19. In contrast, other SP100 isoforms are known to act as transcriptional repressors [43]. With the exception of B2M, these transcript-level changes were not associated with an overall change in gene expression.

Beyond interferon signaling, we observed enrichment of the mRNA translation pathway, with RPS3A exhibiting in T cells a pronounced isoform switch from the shorter RPS3A-208 in controls to the longer RPS3A-201 in COVID-19. This aligns with betacoronavirus-specific splicing patterns previously reported [10], where RPS3A and other ribosomal proteins (RPL10A, RPL5, RPL3) showed differential splicing during SARS-CoV-2 infection. The shift toward RPS3A-201, which encodes the full-length ribosomal protein, suggests enhanced ribosomal biogenesis in infected T cells, potentially facilitating viral hijacking of host translation.

Another notable isoform switch was observed in the anti-apoptotic gene *FAIM*, which protects cells from Fas-mediated apoptosis by blocking the death-inducing signaling complex (DISC) [46]. Prior studies have highlighted the importance of Fas-mediated apoptosis in COVID-19, linking FasL expression to disease severity, inflammation, and survival [47–49]. In ciliated cells, we observe a switch from the longer *FAIM-L* isoform in controls to the shorter, ubiquitously expressed *FAIM-S* isoform in COVID-19, while overall *FAIM* transcript levels remain unchanged. Although *FAIM-L* is classically neuron-specific and *FAIM-S* is broadly anti-apoptotic [46,47,50], their relative roles in ciliated epithelial cells remain to be defined. This isoform shift may represent a regulatory mechanism that limits excessive Fas-mediated apoptosis in the airway epithelium during infection.

We also identified multiple genes with differential transcript usage between moderate and critical COVID-19, most notably *FYN -* a non-receptor tyrosine kinase with a critical role in T cell signaling [51,52]. The full-length isoform, *FYN-202*, was upregulated in T cells in critical cases, potentially driving enhanced T cell receptor (TCR) signaling through its canonical interaction with *LCK*, a central kinase in TCR activation [51]. FYN supports LCK in initiating TCR signaling by phosphorylating immunoreceptor tyrosine-based activation motifs (ITAMs) upon antigen recognition (Fig. 6). This phosphorylation enables recruitment and activation of downstream signaling proteins, ultimately driving T cell activation [53]. *FYN*’s SH2 and SH3 domains mediate interactions with signaling partners, while its kinase domain phosphorylates target proteins [54]. The N-terminal region facilitates membrane localization via lipid modifications [54]. We observed differential usage of three *FYN* isoforms lacking functional domains: *FYN-206/207* (kinase domain deletion) and *FYN-229* (membrane-anchoring domain truncation). These truncated isoforms may act as dominant-negative regulators by competing for interaction partners such as *LCK* but, lacking catalytic activity, they fail to propagate downstream signaling. In contrast, the upregulation of the full-length isoform, *FYN-202,* in critical COVID-19 is consistent with enhanced TCR signal transduction and the hyperactivated T cell state observed in severe disease [55].

**Figure 6:**
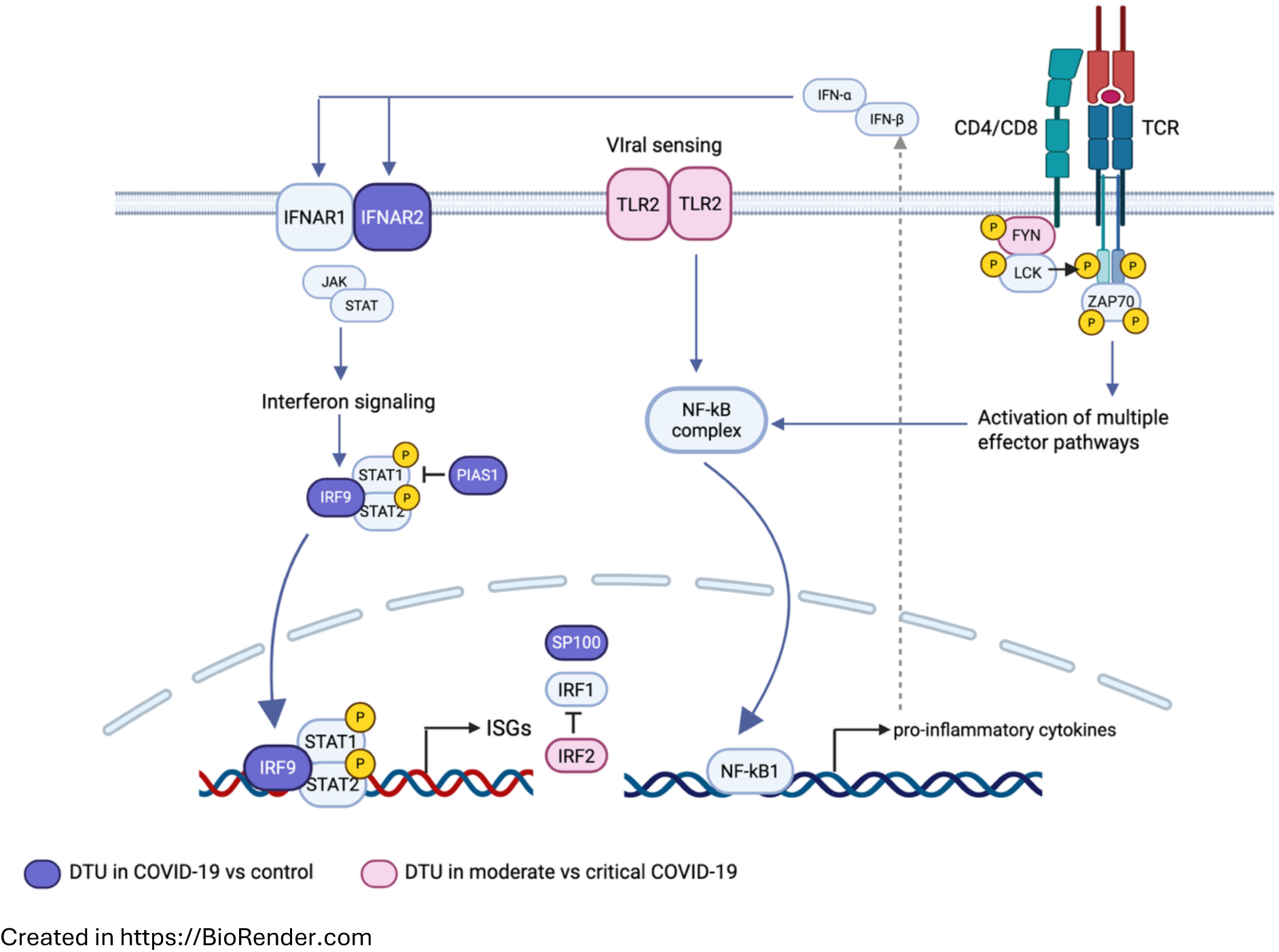
Summary of major pathways regulated by DTU genes in COVID-19. DTU genes identified in moderate vs. critical COVID-19 are highlighted in pink; DTU genes from COVID-19 vs. healthy controls are shown in dark blue. TCR signaling pathway adapted from West et al., 2022; interferon signaling adapted from Bucciol and Meyts, 2023 [36,37]

Pathway analysis of genes with differential transcript usage further highlighted immune response signatures, including enrichment in the Hallmark Interferon Alpha Response and GO Defence Response to Virus pathways. Within these pathways, we detected differential expression of *TLR2* and *IRF2* between moderate and critical COVID-19, both driven by distinct isoform usage. For *TLR2*, the observed isoforms shared the same protein-coding sequence but varied in their 5′ UTRs. *TLR2* can homodimerize and heterodimerize with multiple TLRs and was shown to have both pro- and anti-inflammatory capacities [56,57] and to be associated with COVID-19 disease severity in targeted gene expression profiling [58]. Interestingly, we also detected differential transcript usage for *IRF2*, with isoforms that differed in their 5′ UTRs, suggesting a common mechanism of post-transcriptional regulation for *TLR2* and *IRF2*. *IRF2* encodes a transcription factor that competes with *IRF1* in regulating type I interferon genes, playing a modulatory role in antiviral immunity [59–61]. Variation in 5′ UTRs can influence mRNA stability, translation efficiency, and subcellular localization, offering a way to fine-tune gene expression. Regulation through 5’UTRs results in tight regulation of mRNA and protein levels [62], particularly for genes where changes in dosage are deleterious and lead to disease [63] and may therefore be a strategy for modulation of the immune response in critical COVID-19 via genes like *TLR2* and *IRF2*.

The automated analysis pipeline used in this study retains only known isoforms. As a result, novel or unannotated isoforms that lack Ensembl annotations were systematically filtered out and thus not considered in our analysis. While this approach ensured a conservative and reference-guided analysis, it also limited our ability to explore novel isoforms, which remains an interesting avenue for future investigation. Recent improvements in Nanopore sequencing chemistry and basecalling will enable more accurate splice junction and isoform detection, facilitating both alternative splicing characterization and novel isoform discovery. However, an extensive benchmarking study of long-read RNA-seq methods reported only moderate agreement between bioinformatics tools for transcript identification and quantification [64]. This highlights ongoing challenges in the field, which are further complicated by the lack of definitive ground truth and the need for experimental validation of novel isoforms.

Other limitations of our study include the relatively small number of patient samples with moderate COVID-19, sparsity in the transcript matrices data, and inter-patient heterogeneity. These factors reduced statistical power, particularly for lowly expressed genes, and likely led to under detection of differentially used transcripts. Increased patient numbers and sequencing depth will be important for refining transcript-level insights.

Taken together, our results highlight the power of long-read single-cell sequencing to uncover potential regulatory mechanisms driven by alternative splicing, providing key insights into COVID-19 pathogenesis and immune modulation, and paving the way for future investigations into virus-induced splicing alterations.

## Methods

### Patient Cohort

COVID-19 samples were collected from patients enrolled in the prospective observational cohort study Pa-COVID-19 at Charité Universitätsmedizin Berlin [65]. Human airway specimens were acquired using nasopharyngeal swabs (NSs), as previously described [2], from patients with COVID-19 who were hospitalized at Charité between March and May 2020. Unfragmented single-cell cDNA libraries from seven patients from the cohort were included in this study. Based on WHO guidelines, four of the patients were classified as critical and three as moderate COVID-19 cases (Supplementary Table 2). Further information about the patient’s symptoms, hospitalization and treatment was previously published [2]. Nasal swabs of three controls (C133, C137, C21) who tested negative for SARS-CoV-2 and had no cold-or flu-like symptoms were included. C133 and C137 were recruited for the RECAST study and collected as previously described [26]. Written informed consent was given by all patients and/or their parents before inclusion. Both studies were conducted in accordance with the Declaration of Helsinki and approved by the respective Institutional Review Boards (Pa-COVID-19/RECAST: EA2/066/20). C21 sample was obtained from a consenting laboratory member under approved ethical guidelines.

### Library preparation and short-read sequencing

Single-cell suspensions were prepared from freshly procured human airway specimens. The cell suspensions were converted into barcoded single-cell RNA-seq libraries using the 10X Genomics Single Cell 3 Ĺibrary Kit v3.1 (Cat. No. 1000268) and 10X Chromium Controller, as described previously [2,26]. Aliquots of the amplified cDNA libraries were used for short-read sequencing library preparation following the 10x Genomics protocol and sequenced on a NovaSeq 6000 Sequencer with either S2 or S4 flow cells (Illumina; paired end, single indexing).

### Long-read sequencing

Aliquots from the remaining 10x single-cell cDNA libraries were prepared for nanopore sequencing. Since amplified cDNA libraries contain up to 50% artefactual molecules i.e., cDNA molecules without polyA and cell barcodes, depletion of those artefacts using biotin tags was performed as described in Lebrigand, K., Magnone, V., Barbry, P. et al. [15] to increase sequencing throughput. Briefly, through 5 PCR cycles with 10 ng of the 10x Genomics PCR product, biotin tags were added to the 5’ ends of the cDNA molecules. After removing excess biotinylated primers using AMPure XP Beads (Beckman Coulter) purification, streptavidin beads were utilized to capture the biotinylated full-length transcripts. After removing artefactual molecules by washing the beads, the enriched libraries were re-amplified in 8 cycles to generate sufficient material for long-read library preparation. Library preparation for the control and COVID4 to COVID12 samples was performed using the Oxford Nanopore ligation sequencing kit SQK-LSK112 according to the manufacturer’s protocol. Samples COVID1, 2 and 3 were processed using Oxford Nanopore kit SQK-LSK114. All libraries were sequenced on a PromethION24 sequencer (Oxford Nanopore). Base-calling was performed using the super accurate model in guppy, Version 6.4.6, using the configuration file dna r10.4 e8.1 sup.cfg for kit 12 samples and dna r10.4.1 e8.2 260bps sup.cfg for kit 14 samples.

### Short-read data processing

Pre-processing of the raw short-read data was performed as described before [2,26]. The analysis in those studies was carried out with Seurat version 3.1.4. Subsequent analyses were performed in Seurat v5.1.0 by upgrading the existing Seurat objects. Expression values were log-normalized, variance-scaled, and centered using `NormalizeData` and `ScaleData`. Highly variable genes were identified with `FindVariableFeatures`, and principal component analysis was applied via *`RunPCA`* to compute the top 50 PCs. Batch effects were corrected using Harmony v1.2.1 on these PCs. A shared k-nearest-neighbor graph was constructed with `FindNeighbors`, clusters were identified using `FindClusters`, and cells were projected into two dimensions with `RunUMAP` (first 30 PCs). In iterative downstream steps, clusters were manually inspected for quality and annotated based on epithelial and immune marker expression, with clustering resolution adjusted as needed to refine cell-type assignments.

### Long-read data processing

Long-read data were processed with the `epi2me-labs/wf-single-cell` workflow [66], version 2.0.2, employing kit-specific configurations and default parameters [66]. Reads were aligned to the custom reference genome comprising both the human and SARS-CoV-2 sequences. The workflow extracts and corrects cell barcodes and UMIs, then assigns each read to its corresponding gene and annotated transcript isoform. Output files mirror those produced by CellRanger, yielding both gene-by-cell and transcript-by-cell expression matrices.

Gene expression matrices were imported into Seurat, version 5.1.0 [67]. Cells expressing fewer than 200 genes or more than 15 % mitochondrial reads were excluded. Sample-specific upper UMI count thresholds (10,000 for C133 to 100,000 for COVID1) were applied to mitigate doublet artifacts. An assay containing transcript-level counts was added, but all integration and dimensionality-reduction steps were performed on the gene-expression assay due to the high sparsity of transcript-level data. Normalization, scaling, integration, dimensionality reduction and annotation were performed as described above for the short-read data. Additionally, transcript-level counts were log-normalized to stabilize variance for downstream visualization.

Transcript biotype annotations were downloaded from Ensembl BioMart (GRCh38.p14) [68].

### Statistical analysis

Immune and epithelial cell counts were compared between control and COVID-19 samples using a two-sided Fisher’s exact test on the K×2 contingency table of cell-type versus condition. For each immune cell type, PTPRC isoform expression was compared against all other immune cells using Seurat’s `FindMarkers (test.use =` “`wilcox`”), which performs a two-sided Wilcoxon rank-sum test on normalized counts.

### Comparison of short- and long-read data

To benchmark nanopore–based single-cell RNA-Seq against the standard short-read approach, we split the same cDNA library into two aliquots and generated long- and short-read data from identical cells. Cell barcodes were used to link per-cell statistics and annotations across both modalities. We applied Seurat’s `FindTransferAnchors` function in the PCA space of the short-read dataset, followed by `TransferData` to project cell-type labels onto the long-read dataset. A transfer score is computed for each cell in the long-read dataset, reflecting the confidence of the assigned label based on the similarity of its PCA-transformed expression profile to that of reference cells in the short-read dataset. We then computed the average transfer score for each predicted label to quantify the overall concordance between cells of the same type in each dataset.

### Differential gene expression and transcript usage testing

For differential gene expression between controls and COVID-19 severity groups, we aggregated (pseudo-bulk) gene counts per cell type and analyzed them with DESeq2, version 1.42.0 [31]. Only genes with ≥10 total counts in at least three samples (the size of the smallest comparison group) were tested.

To detect changes in isoform usage, we applied DEXSeq [32], version 1.48.0, to aggregated transcript counts per cell type. We restricted testing to genes expressed in at least nine samples for the control vs. COVID-19 comparison (or six samples for moderate vs. critical) and encoding at least two annotated isoforms. Within each gene, individual isoforms were required to have ≥5 counts in at least two-thirds of the samples in the smaller group to be included in the analysis.

### Gene set enrichment analysis

Genes tested by DESeq2 were ranked by their Wald statistic, which combines log₂ fold-change magnitude and standard error. Hallmark, Reactome, and GO Biological Process gene sets [35] were downloaded from MSigDB release 2024.1.Hs [34]. Reactome pathway enrichment was performed with `gsePathway` (ReactomePA, version 1.48.0) [69,70] using 100 000 permutations. GO and Hallmark enrichments were computed via `GSEA` (clusterProfiler, version 4.12.6) [71]. For each cell type, the top eight Reactome categories (ordered by adjusted p-value) were extracted, annotated by function, and displayed in a combined heatmap. The results were filtered for readability, more general categories (Interferon Signaling, NMD) were removed when the results contained more specific categories, e.g. Interferon alpha/beta signaling, Interferon Gamma signaling with similar enrichment scores. Enrichment plots were generated with `gseaplot2` (enrichplot, version 1.24.4) [72] and then customized in ggplot2, version 3.5.1, to include an additional track highlighting genes with differential transcript usage.

## Data availability

Raw long-read sequencing data are deposited in the European Genome-phenome Archive (EGA) under controlled access (accession EGAS50000001290). Access requires a Data Transfer Agreement. Metadata tables (patient ID, severity, sex, age), cell count matrices for both short- and long-read data, and differential testing results are available via <u>Zenodo</u>.

## Acknowledgements

We thank all patients of the RECAST and Pa-COVID-19 studies for donating nasal and nasopharyngeal samples and clinical data. This project has received funding from EASI-Genomics, the European Union’s Horizon 2020 research and innovation program under grant agreement No 824110. The study was supported by the Medical Scientist Program of Charité Universitätsmedizin-Berlin, the Alliance4Rare Network of the Eva Luise and Horst Köhler Stiftung and the Alexander von Humboldt Foundation. Additionally, we would like to thank Dr. Marco Binder (Dynamics of Early Viral Infection and the Innate Antiviral Response, Center for Integrative Infectious Diseases Research, CIID, University Heidelberg) for his expertise in viral infections and his valuable input on this study. The authors declare no competing interests.

## Author Contributions

P.N.R., I.L., R.E. and C.C. acquired funding and supervised the work. A.R. conceived and designed the experiments. K.K. designed the data processing pipeline and implemented it with support from L.M.. K.K., A.R. and L.M. conducted all computational analyses. K.K. and L.M. interpreted the data. A.R. and M.B. performed long-read sequencing library preparation and sequencing. R.L.C. and J.L. performed 10x Genomics sample processing as well as short-read library preparation. K.K., L.M. and A.R. wrote the manuscript with contributions from all authors. All authors read and approved the final manuscript.

## Declaration of interests

The authors declare no competing interests.

## Supplements

**Figure S1:**
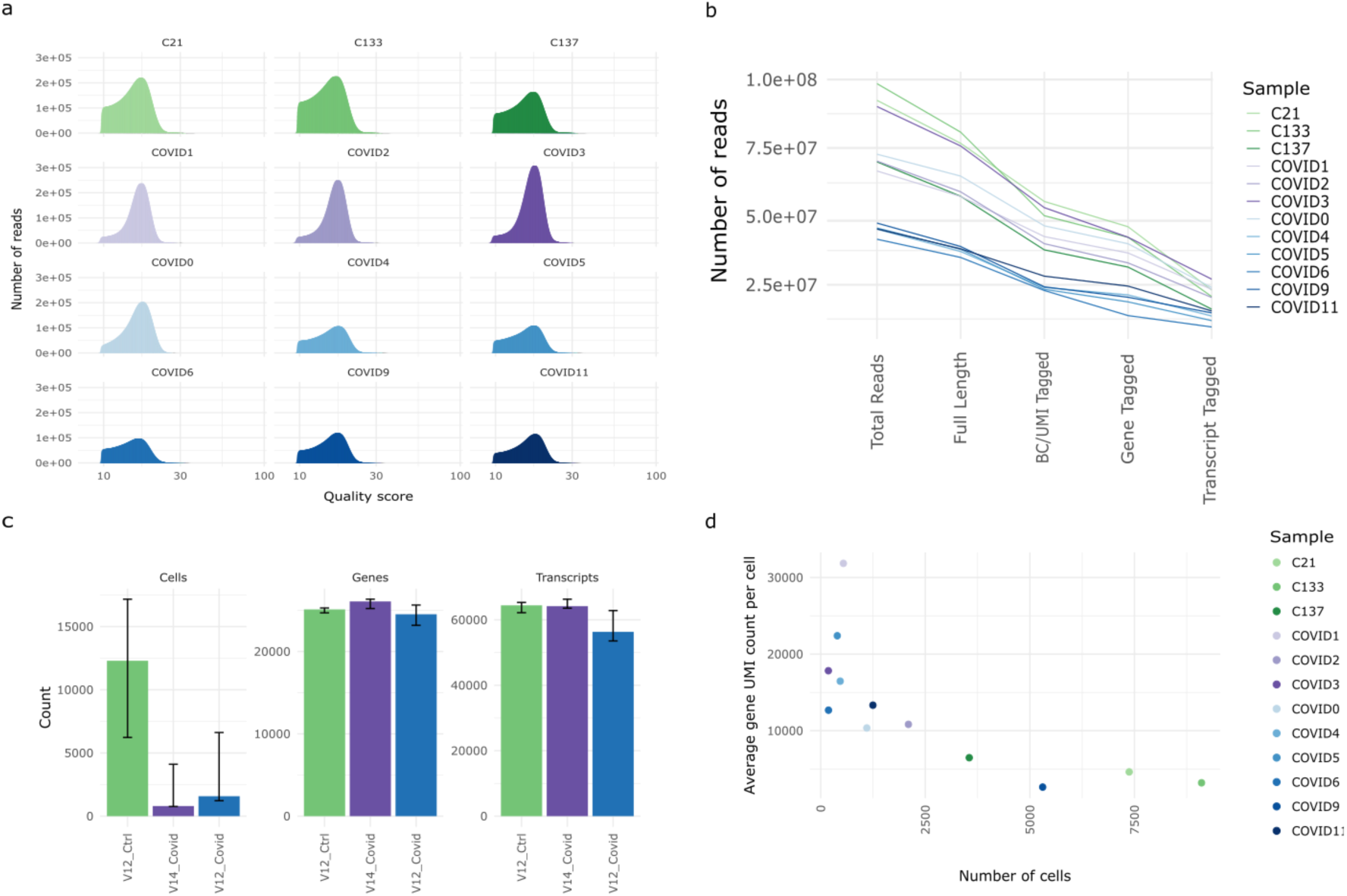
(a) PHRED scores of reads that passed QC, samples sequenced with V14chemistry (purple) exhibit higher read quality (b) wf-single-cell read statistics: counts of reads with the expected full-length configuration (correct 5′ and 3′ adapters), reads successfully assigned a valid barcode + UMI, and reads mapped to a known gene and isoform (c) Median number of cells, detected genes and transcript isoforms across conditions and chemistries, error bars indicating total range of values (d) Relationship between average UMI count per cell and cell number per sample (after QC in Seurat).

**Figure S2:**
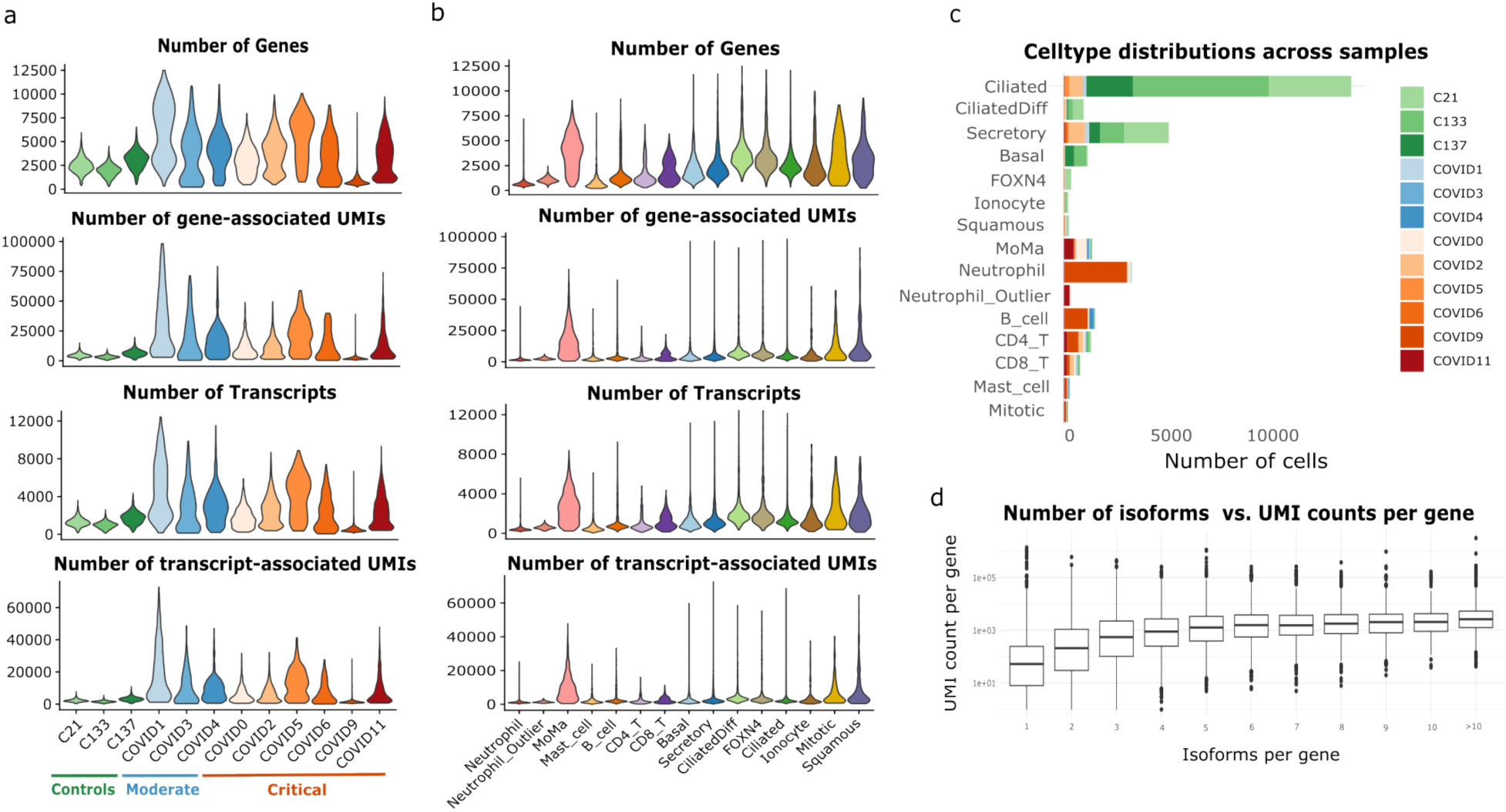
(a) Number of detected genes, transcripts and corresponding UMI counts per sample; control samples exhibit lower numbers of per-cell counts due to increased number of cell (b) Number of detected genes, transcripts and corresponding UMI counts per cell type; epithelial cell types and Monocytes/Macrophages exhibit more variable gene and transcript expression (c) Distribution of cell types across samples; COVID-19 samples are characterized by immune cell infiltration (d) Relationship between the number of isoforms detected per gene and sequencing depth (UMI count per gene)

**Figure S3:**
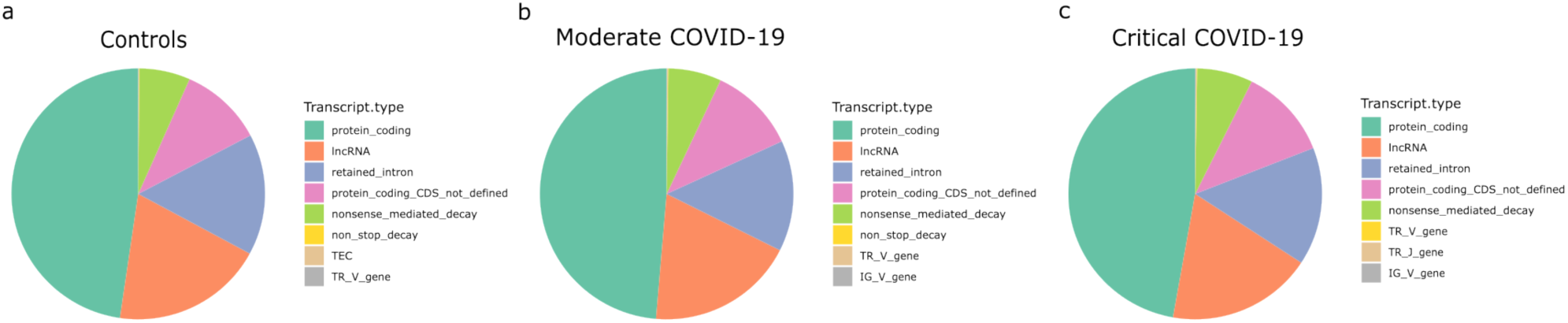
Pie charts depicting the proportions of isoforms identified by wf-single-cell and categorized by Ensembl biotype for (a) isoforms identified in control samples (b) isoforms identified in moderate COVID-19 (c) isoforms identified in critical COVID-19

**Figure S4:**
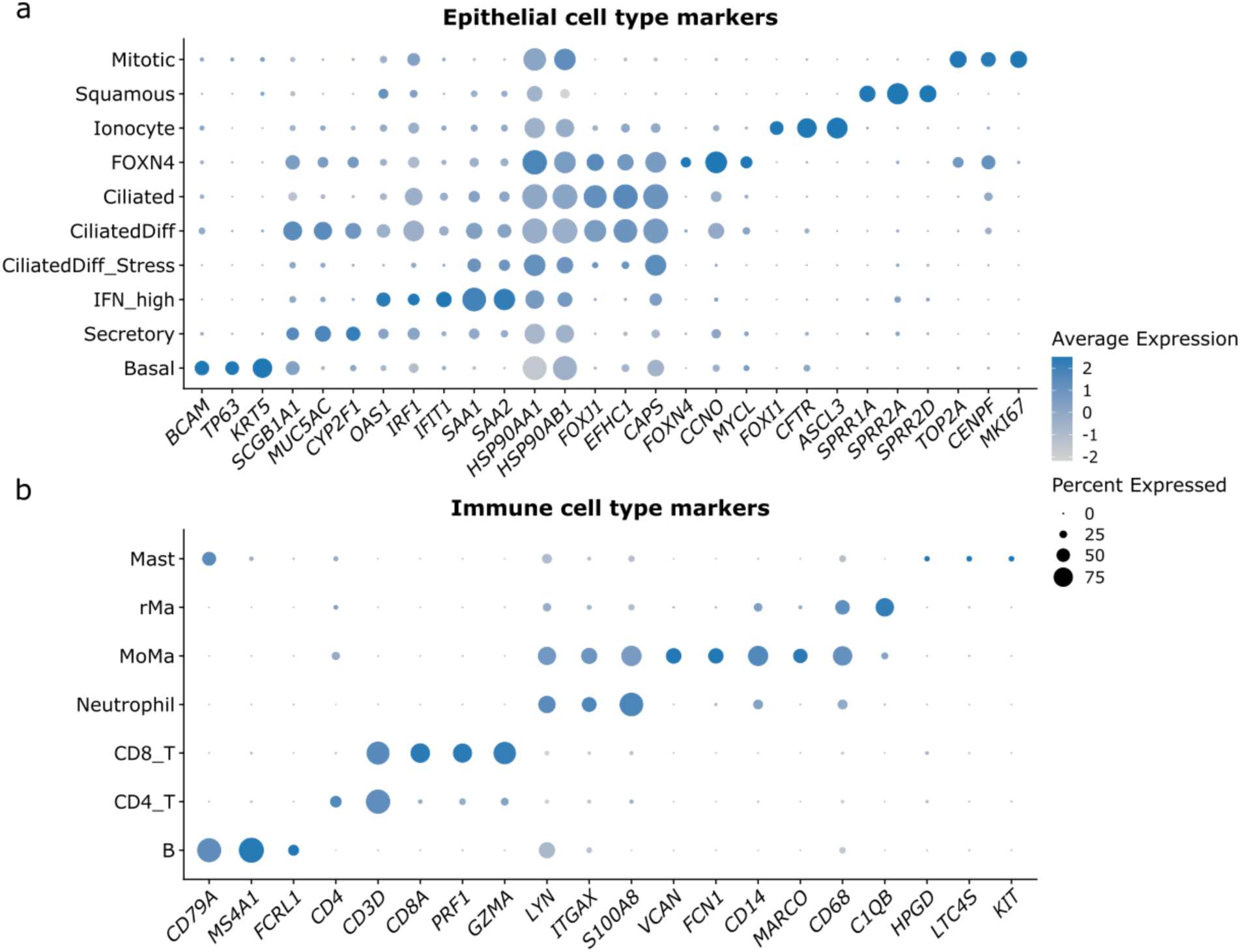
(a) Dot plots displaying marker gene expression for epithelial cell populations identified in the short-read data (b) Dot plots displaying marker gene expression for immune cell populations identified in the short-read data

**Figure S5:**
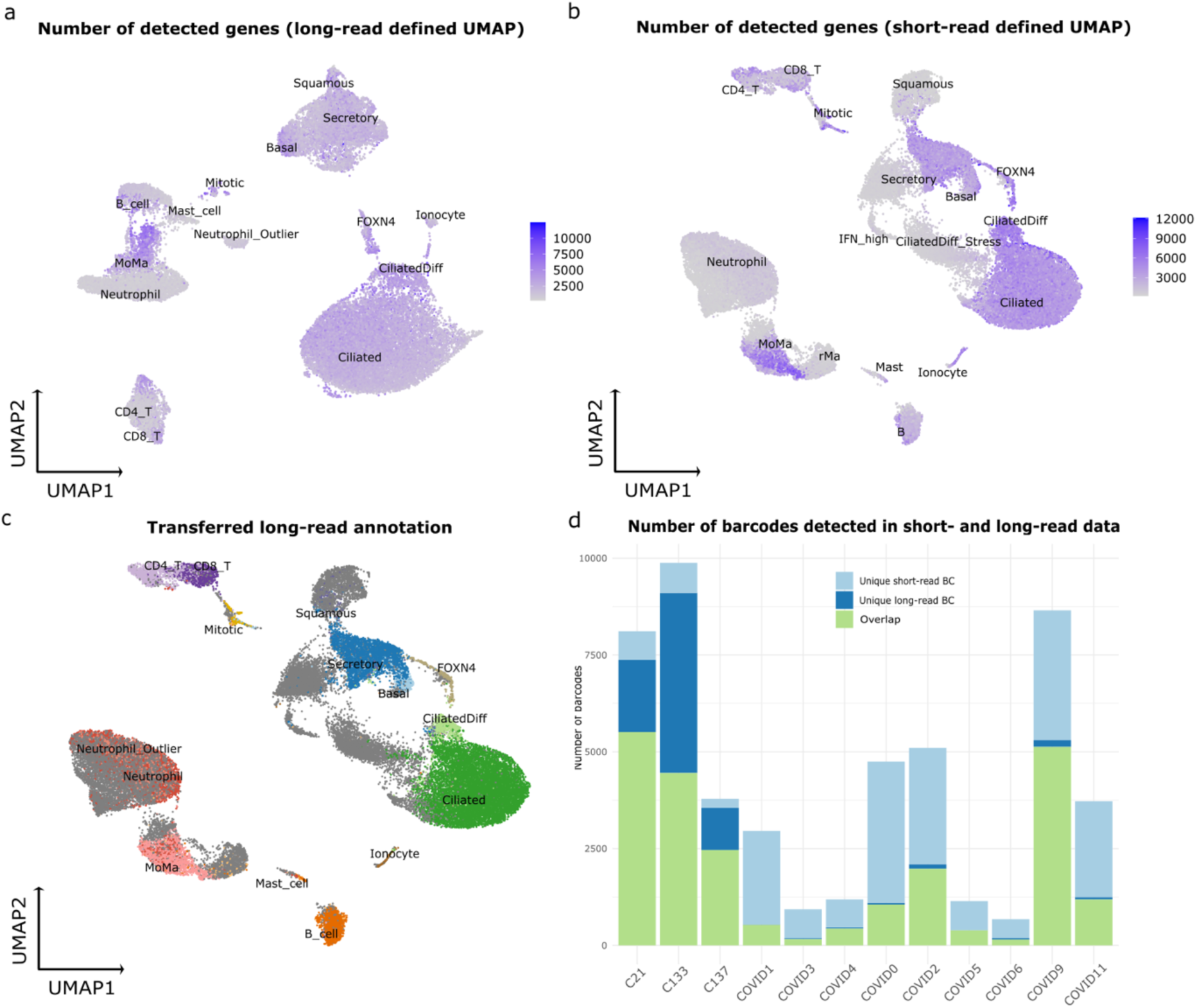
(a) UMAP of gene expression from short-read data: cell types with greater complexity show more detected genes (b) UMAP of gene expression from long-read data: again, more complex cell types show higher gene counts (c) Long-read cell annotations projected onto the short-read UMAP: cell populations with low gene detection in the short-read data are missing in the long-read dataset (d) Comparison of cell barcodes between short- and long-read data: long-read sequencing detects more cells in control samples, whereas COVID-19–derived cells are better retained in the short-read data

**Figure S6:**
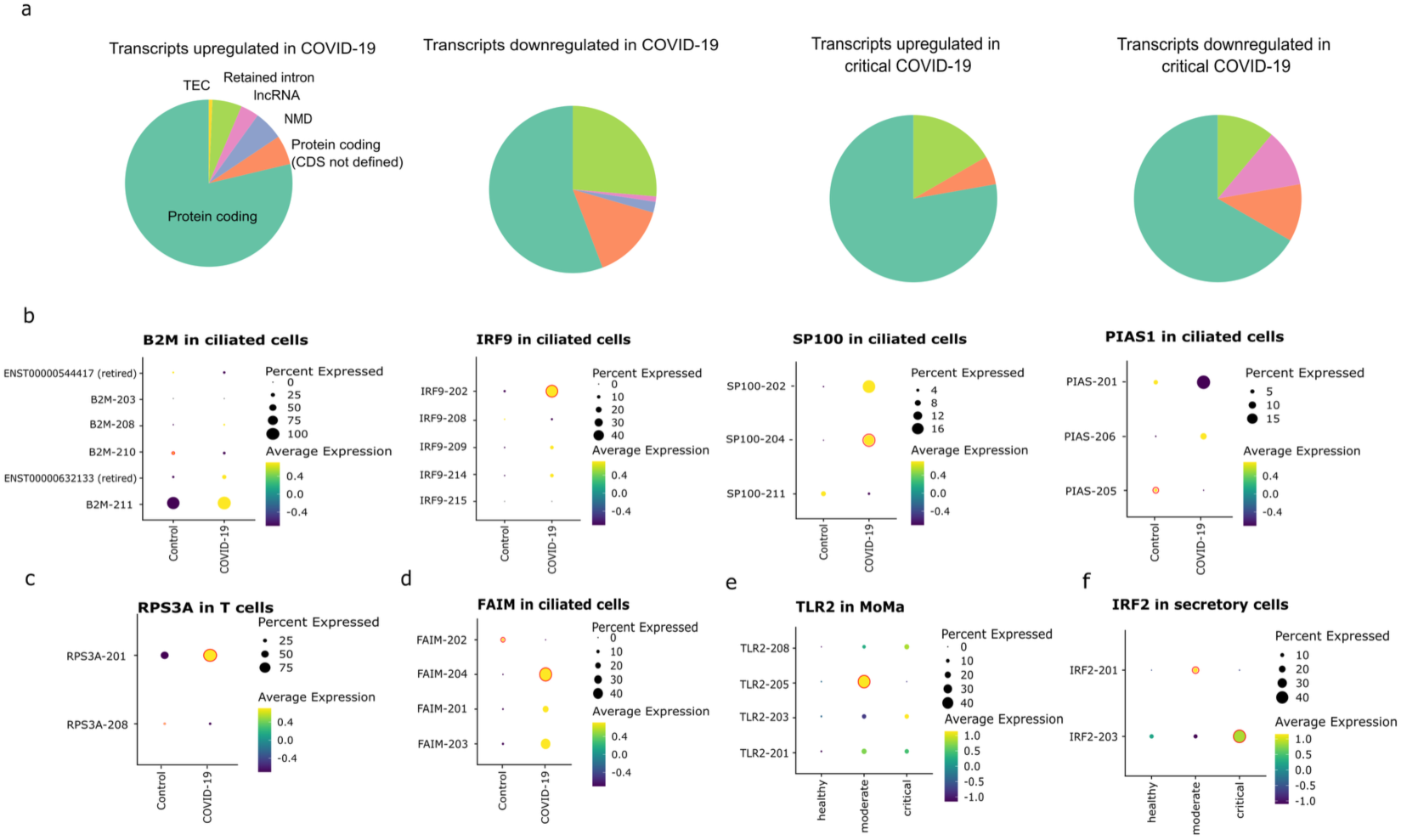
(a) Pie charts displaying the biotypes of transcripts identified as differentially used between control vs. COVID-19 (left) and moderate vs. critical COVID-19 (right). (b) Dot plots showing genes with DTU in ciliated cells that are involved in *Interferon gamma signaling (B2M-210:* log_2_FC = −0.68, p_adj_ = 0.02, *IRF9:* log_2_FC = 1.31, p_adj_ = 0.07, *SP100:* log_2_FC = 0.51, p_adj_ = 0.05, *PIAS1-205:* log_2_FC = −3.02, p_adj_ = 1.9 x 10^-3^). (c) Dot plot showing expression of *RPS3A* isoforms across T cells. A significant change in transcript usage is observed between conditions, with a shift from *RPS3A-208* to *RPS3A-201* (log_2_FC = −1.21, p_adj_ = 0.04 for *RPS3A-208*; log_2_FC = 0.56, p_adj_ = 0.03 for *RPS3A-201*). (d) Dot plot showing expression of *FAIM* isoforms in ciliated cells. A significant change in transcript usage is observed between conditions, with a shift from *FAIM-202* to *FAIM-204* (log_2_FC = - 19.31, p_adj_ = 0.01for *FAIM-202*; log_2_FC = 2.05, p_adj_ = 3.7 x 10^-8^ for *FAIM-204*). (e) Dot plot showing expression of *TLR2* isoforms across Monocytes and Macrophages. A significant decrease in transcript usage of TLR2-205 is observed in critical COVID-19 (log_2_FC = −20.60, p_adj_ = 0.07). (f) Dot plot showing expression of *IRF2* isoforms across secretory cells. We observe a significant shift from *IRF2-201* to IRF2-203 isoform usage in critical COVID-19 (log_2_FC = −4.31, p_adj_ = 1.5 x 10^-5^ for *IRF2-201*; log_2_FC = 1.41, p_adj_ = 3.4 x 10^-6^ for *IRF2-203*)

**Table S1:** Clinical information about samples used for this study.

| Sample No. | Sample | WHO Categorization | Age | Sex |
| --- | --- | --- | --- | --- |
| COVID0 | Nasopharynx | critical | 63 | m |
| COVID1 | Nasopharynx | moderate | 71 | m |
| COVID2 | Nasopharynx | critical | 32 | m |
| COVID3 | Nasopharynx | moderate | 52 | m |
| COVID4 | Nasopharynx | moderate | 64 | m |
| COVID5 | Nasopharynx | critical | 54 | m |
| COVID6 | Nasopharynx | critical | 62 | f |
| COVID9 | Nasopharynx | critical | 50 | f |
| COVID11 | Nasopharynx | critical | 73 | m |
| C21 | Nasopharynx | control | 29 | m |
| C133 | Nasal | control | 66 | m |
| C137 | Nasal | control | 62 | f |

**Table S2:**
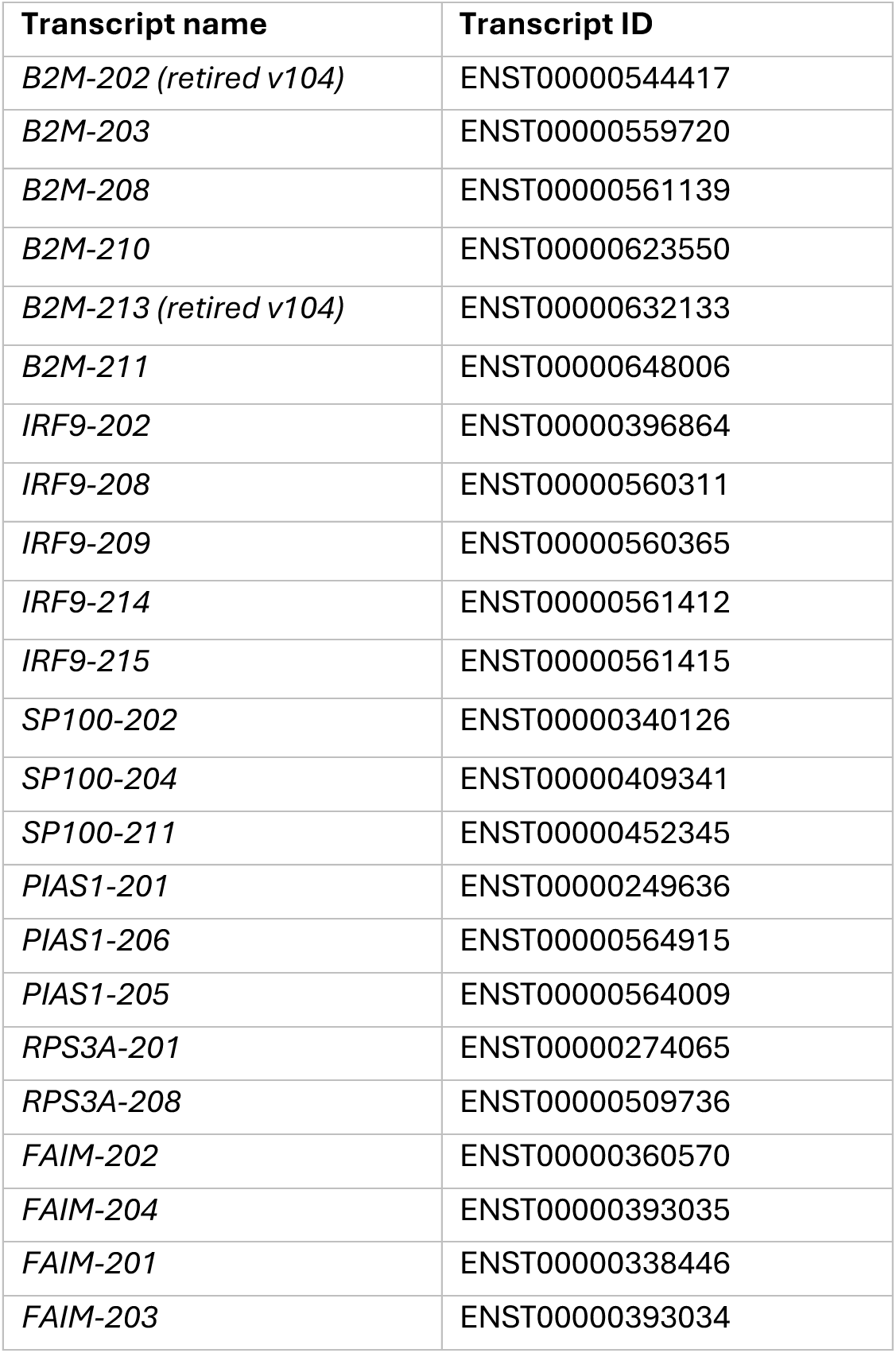
Ensembl transcript names and identifiers of genes with DTU between control and COVID-19.

| Transcript name | Transcript ID |
| --- | --- |
| *B2M-202 (retired v104)* | ENST00000544417 |
| *B2M-203* | ENST00000559720 |
| *B2M-208* | ENST00000561139 |
| *B2M-210* | ENST00000623550 |
| *B2M-213 (retired v104)* | ENST00000632133 |
| *B2M-211* | ENST00000648006 |
| *IRF9-202* | ENST00000396864 |
| *IRF9-208* | ENST00000560311 |
| *IRF9-209* | ENST00000560365 |
| *IRF9-214* | ENST00000561412 |
| *IRF9-215* | ENST00000561415 |
| *SP100-202* | ENST00000340126 |
| *SP100-204* | ENST00000409341 |
| *SP100-211* | ENST00000452345 |
| *PIAS1-201* | ENST00000249636 |
| *PIAS1-206* | ENST00000564915 |
| *PIAS1-205* | ENST00000564009 |
| *RPS3A-201* | ENST00000274065 |
| *RPS3A-208* | ENST00000509736 |
| *FAIM-202* | ENST00000360570 |
| *FAIM-204* | ENST00000393035 |
| *FAIM-201* | ENST00000338446 |
| *FAIM-203* | ENST00000393034 |

**Table S3:** Ensembl transcript names and identifiers of genes with DTU between moderate and critical COVID-19.

| Transcript name | Transcript ID |
| --- | --- |
| *FYN-229* | ENST00000538466 |
| *FYN-207* | ENST00000462856 |
| *FYN-206* | ENST00000462598 |
| *FYN-202* | ENST00000354650 |
| *IRF2-201* | ENST00000393593 |
| *IRF2-203* | ENST00000504340 |
| *TLR2-208* | ENST00000646900 |
| *TLR2-205* | ENST00000643501 |
| *TLR2-203* | ENST00000642700 |
| *TLR2-201* | ENST00000260010 |

## Notes

### Competing Interest Statement

The authors have declared no competing interest.

